## Supplementary material for "Basal Contamination of Sequencing: Lessons from the GTEx dataset": Labels for Supplemental Tables

### Description of Additional Supplementary Files

**File Name: Supplementary Data 1**

**Description:** Ratio of tissues sequenced on pancreas and non-pancreas days.

**File Name: Supplementary Data 2**

**Description:** Ratio of tissues sequenced on esophagus and non-esophagus days.

**File Name: Supplementary Data 3**

**Description:** The output of the pancreas gene contamination linear mixed model.

**File Name: Supplementary Data 4**

**Description:** The output of the esophagus mucosa gene contamination linear mixed model.

**File Name: Supplementary Data 5**

**Description:** A technical comparison of the GTEX1 fibroblast sample and its main contaminating GTEX2 esophagus sample.

**File Name: Supplementary Data 6**

**Description:** Ten RNA-seq experiments with two or more tissues/cells all demonstrating cross contamination between samples.
